# A gelatin-based feed for individually tailored drug delivery to adult zebrafish

**DOI:** 10.1101/2022.10.11.511627

**Authors:** Aleksander J. Ochocki, Justin W. Kenney

## Abstract

Current approaches to drug delivery in adult zebrafish have significant limitations such as need for confinement, anesthesia, and/or dosing that is not based on body weight. To circumvent these challenges, we developed a non-invasive gelatin-based feed that is easily pipette into individually tailored morsels according to weight. Our feed was readily eaten by zebrafish (< 1 minute) with minimal leaching of compound (< 5%) when placed in water. We used our feed to deliver an NMDAR antagonist (MK-801, 4 mg/kg) prior to exposure to a novel tank. Consistent with prior work, we found that MK-801 caused a decrease in predator-avoidance/anxiety-like behaviors. We also found that MK-801 increased locomotion in male fish, but not females. Our simple, easy to prepare, and individually tailored gelatin-based feed now brings pharmacological manipulations of adult zebrafish in line with best practices used in other vertebrate model organisms.

## Introduction

Zebrafish were first suggested as a model organism in embryology nearly a century ago due to their ease of maintenance, fecundity, and transparent embryos (Creaser, 1934). In the 1980’s, they finally became established as a model in developmental biology (Grunwald and Eisen, 2002), and near the turn of the 21^st^ century zebrafish were taken up more broadly in fields like neuroscience, immunology, and regenerative medicine (Kenney, 2020; Norton and Bally-Cuif, 2010; Poss et al., 2003; Trede et al., 2004). This increase in uptake was driven by the development of a sophisticated genetic toolbox and a deeper appreciation for the insight zebrafish provide into vertebrate evolution and biology. The ascent of zebrafish stands to continue thanks to the creation of advanced digital resources for studying complex organ systems like the brain (Kenney et al., 2021; Kunst et al., 2019; Randlett et al., 2015; Ronneberger et al., 2012; Tabor et al., 2019) and vasculature (Kugler et al., 2022). Nonetheless, given that zebrafish are a relative newcomer to the pantheon of model organisms, methodological improvements are still needed to fully realize the potential of zebrafish, particularly at adult stages.

Pharmacological manipulations are an effective approach for identifying molecular contributions to physiological function. Many compounds developed and tested in mammalian systems are often effective in zebrafish, and vice versa, because of the significant overlap between the zebrafish and human genomes (Howe et al., 2013), and the functional and chemical overlap in many physiological systems such as the brain (Panula et al., 2010), immune system (Forn-Cuní et al., 2017), and heart (Vornanen and Hassinen, 2016). This extensive overlap, along with their low cost and early life transparency, has led to the increasing use of zebrafish in drug discovery (MacRae and Peterson, 2015) and early-stage toxicological assessment (Garcia et al., 2016). However, challenges remain for drug delivery to adult animals.

Both adult and larval zebrafish have been used to address various aspects of vertebrate biology, with each stage having its advantages and disadvantages. Larval zebrafish have the benefit of early life transparency, making them widely used in developmental biology. Drug administration in larval animals is also straightforward: compounds are added to water during development with minimal disturbance of animals. Adult stages have the advantages of fully developed organ systems and more sophisticated behaviors, but drug administration presents more of a challenge because fish cannot be confined to the small spaces of multi-well plates or petri dishes like larval animals. Several methods of drug delivery have been developed for adult animals, each with significant drawbacks. The most common method is beaker or tank dosing where the drug is dissolved in a large (typically > 100 mL) volume of water and animals are placed in the solution (e.g., Levin et al., 2006; Montgomery et al., 2010; Sison and Gerlai, 2011). The confined space of a beaker is a stressful manipulation which can interfere with the interpretation of experimental results. Drug administration in an entire tank is less stressful but more wasteful as only a small fraction of the drug is absorbed by the animal and it is difficult to know the actual dose received. Additionally, both tank and beaker dosing are also limited to compounds that are available in large quantities. Two injection-based methods have been developed for adult animals: oral gavage and intraperitoneal injections (Collymore et al., 2013; Kinkel et al., 2010). However, both methods require anesthesia and extensive handling. This introduces additional challenges because anesthesia can interact with drug effects and the need for handling reduces throughput and increases the risk of injury to animals.

Delivering drugs via feeding has the potential to be both non-invasive and minimally wasteful. Although a handful of attempts have been made to develop feed-based drug delivery, prior approaches have been hampered by the inability to give precise doses based on body weight, which is the standard of practice in rodent and primate pharmacology. This is because they have relied on either cutting and weighing small amounts of heterogenous food (Sciarra et al., 2014), which is slow and error prone, or the same amount of food is given to all animals based on an average weight (Chang et al., 2017; Lu and Patton, 2022; Zang et al., 2011), relying on the faulty assumption that fish do not vary much in size. To overcome these limitations, we have developed an inexpensive gelatin-based feed method for drug delivery to adult zebrafish that is individually tailored to each animal with ease.

## Results

### Development of gelatin feed

Our goal was to create a feed-based drug delivery system that would allow us to deliver precise amounts of drug to adult zebrafish based on individual body weight. We used gelatin as the base for our feed because of its low melting point and common usage as a food stabilizer. We mixed in spirulina to add color, palatability, and nutrition. When warmed, we were able to pipette precise volumes of feed onto parafilm before solidification at −20 C (Fig. 1A). Because the feed is administered in an aqueous environment where added compound could potentially leach into the water, we modeled drug loss at different time points and formulations using methylene blue dye. Initially, we kept the amount of spirulina constant (4% w/v) and varied gelatin concentration (Fig. 1B). Using a 3 × 4 (gelatin concentration × time) ANOVA, we found a main effect of time (F(3,60) = 173, P < 10^−15^), gelatin (F(2,60) = 4.6, P = 0.014) and an interaction (F(6,60) = 3.63, P = 0.0039). At five minutes, about 5% of methylene blue leached out for all three concentrations of gelatin, but by 10 minutes, more methylene blue leached out of the highest gelatin concentration than the lower concentrations. To determine if spirulina contributed to leaching, we kept the gelatin concentration constant (12% w/v) and varied the concentration of spirulina (Fig. 1C). A 3 × 4 (spirulina concentration × time) ANOVA found a main effect of time (F(3,60) = 60, P < 10^−15^) and spirulina (F(2,60) = 32, P = 2.8 × 10^−10^) with a trend towards an interaction (F(6,60) = 1.9, P = 0.093). Increasing the concentration of spirulina resulted in a decrease in leaching that was evident at each time point. Taken together, we found that altering the concentration of spirulina, but not gelatin, had a clear effect on the amount of methylene blue that leached out of the feed. Based on these data, we used a 12% gelatin, 4% spirulina formulation, noting that less than 5% of the compound leaches from this formulation after 5 minutes.

**Figure 1.**
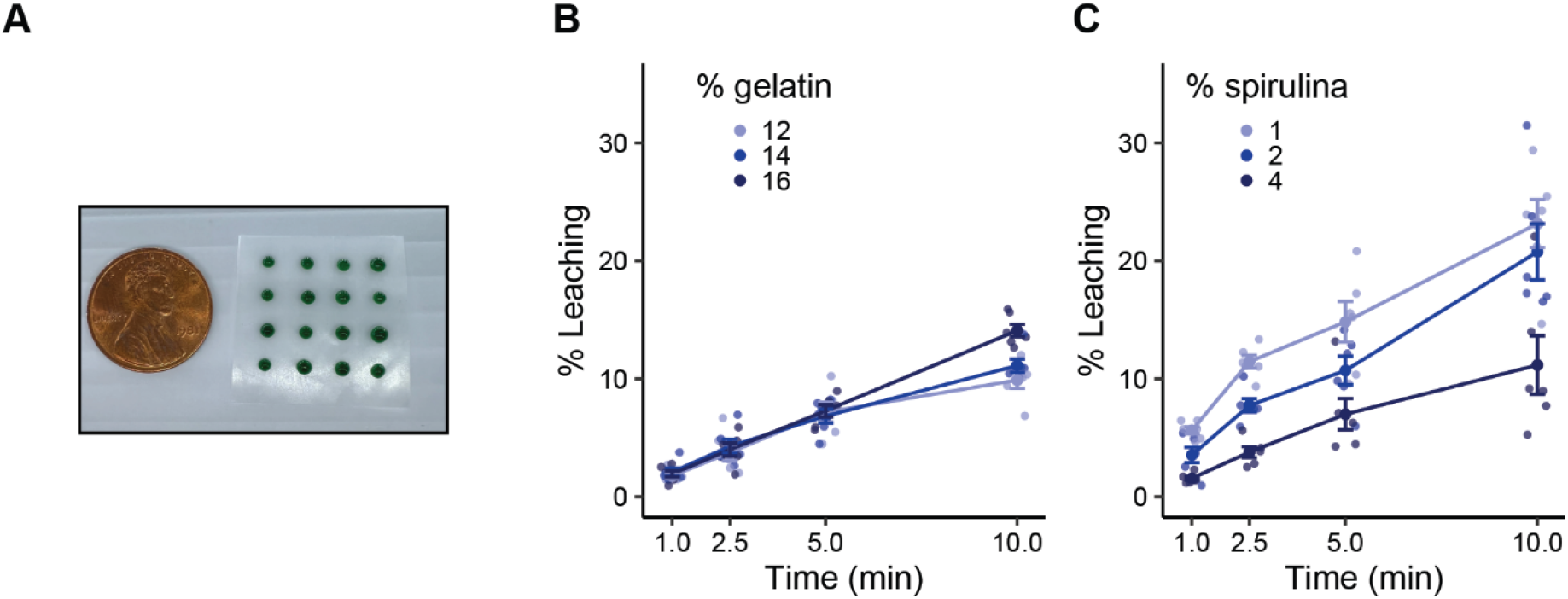
Preparation of gelatin feed and assessment of leaching. A) Gelatin based feed was made by mixing brine shrimp extract, spirulina, and gelatin. While still liquid, gelatin is pipette into individually tailored morsels onto parafilm before setting at −20 °C. B) Methylene blue leaching over time at different gelatin concentrations with 4% w/v spirulina v. C) Methylene blue leaching over time at different spirulina concentrations with 12% w/v gelatin. Data are presented as mean ± SEM, n’s = 6.

### Gelatin feed palatability

Next, we sought to determine if the feed we developed would be readily eaten by adult zebrafish of two commonly used inbred fish strains: ABs and TLs. Fish were given the gelatin feed at 1% body weight for five consecutive days in lieu of their normal morning feed. We found that all (16/16) TL fish ate the feed within five minutes from the very first exposure, but it took 4 consecutive days for 15 of 16 AB fish to consistently consume the feed (Fig. 2A). For time to eat, a 2 × 5 ANOVA (strain × day) found a main effect of strain (F(1,135) = 18.6, P = 3.1 × 10^−5^), a main effect of day (F(4,135) = 5.54, P = 0.00036) and an interaction between strain and day (F(4, 135) = 3.21, P = 0.015). TL fish ate the food quickly from their first exposure (range: 4 – 20 s), improving to under 10 s for all fish by the fourth day. AB fish took longer to eat initially (day 1 range: 7 – 144 s), but by the fourth and fifth days all fish, except one, ate within 10 s. Thus, once acclimated, fish typically ate the gelatin feed in under 30 seconds, which is well before any appreciable drug leaching could occur (Fig. 1).

**Figure 2.**
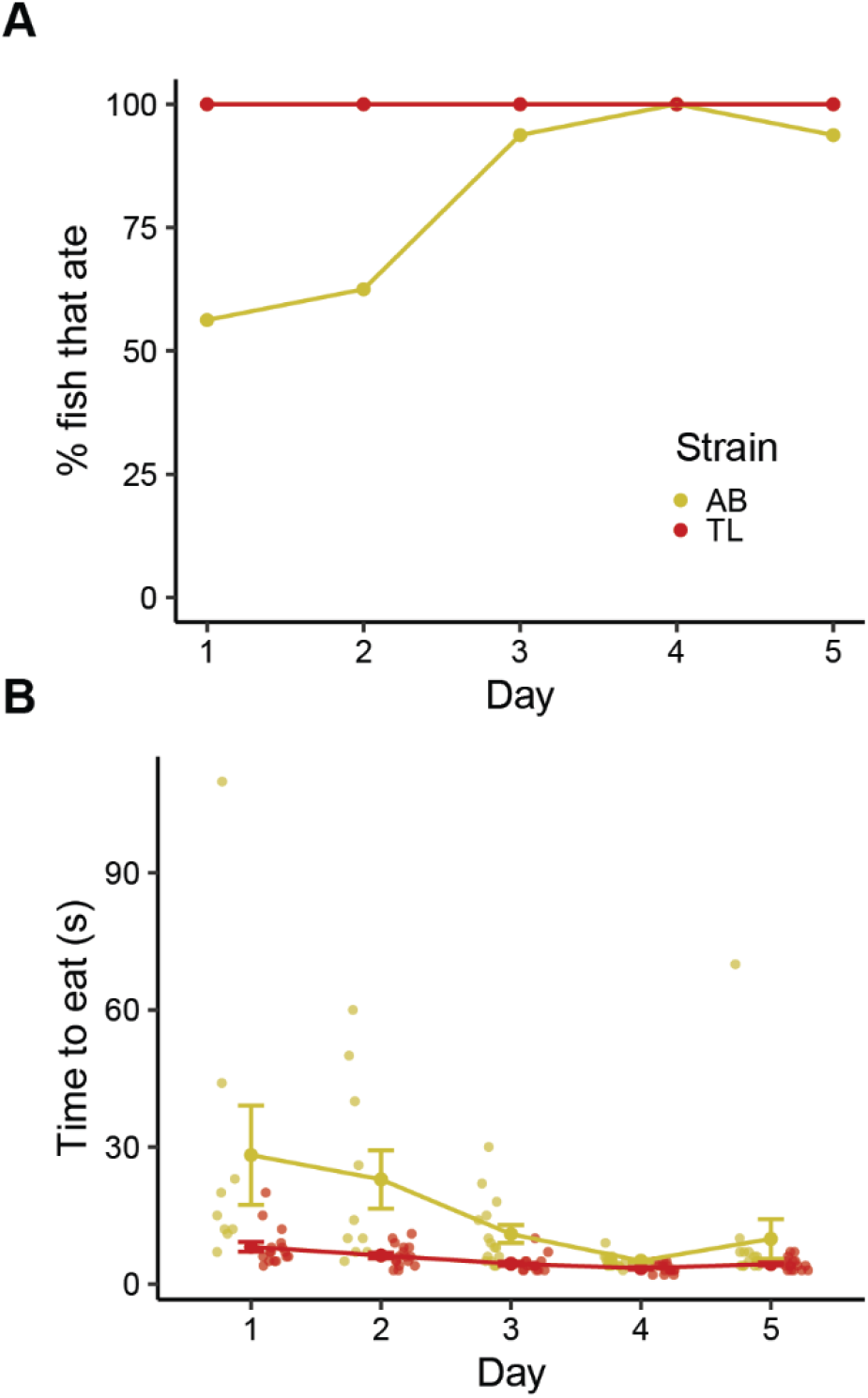
Eating of gelatin-based feed by fish. A) Percent of fish from two strains that ate the feed within five minutes of administration. B) Time it took for AB or TL fish to eat the gelatin feed. Fish that did not eat the feed were excluded from analysis. Data presented as mean ± SEM, n = 16 fish per strain.

### Behavioral effect of MK-801 administered using gelatin feed

As a proof-of-principle to determine if our gelatin-based feed could successfully deliver a drug that is known to affect behavior, we gave MK-801, a glutamatergic NMDAR (N-methyl-D-aspartate receptor) antagonist, to AB fish and measured their exploration of a novel tank. MK-801 has been widely used in adult zebrafish and is known to affect predator avoidance/anxiety-like behaviors, memory, and locomotion (Kenney et al., 2017; Seibt et al., 2011; Sison and Gerlai, 2011). Fish were given either vehicle or 4 mg/kg of MK-801 thirty minutes prior to being placed in a novel tank for six minutes where we measured geotaxis (bottom distance), thigmotaxis (center distance), and distance travelled (Fig. 3). We used 2 × 2 (sex × drug) ANOVAs to assess statistical significance. For bottom distance (Fig. 3A) we found a main effect of drug (F(1,69) = 24.5, P = 0.000051) where fish given MK-801 spent more time near the top of the tank. There was no effect of sex (F(1, 69) = 2.52, P = 0.12) or an interaction (F(1,69) = 1.0, P = 0.32). Post-hoc FDR corrected t-tests within sex found that MK-801 increased bottom distance in both female (P = 0.0076) and male (P = 0.023) fish. For center distance, there was a trend towards an effect of drug (F(1,69) = 3.04, P = 0.084) where fish given MK-801 appeared to spend more time near the center of the tank. There was an effect of sex (F(1,69) = 5.84, P = 0.014), but no interaction (F(1,69) = 0.76, P = 0.76) where males, irrespective of treatment, were closer to the center of the tank than females. However, FDR corrected post-hoc t-tests found no effect of drug in either female (P = 0.35) or male (P = 0.25) animals. Finally, for distance travelled, there was a main effect of drug (F(1,69) = 4.2, P = 0.044), with fish given MK-801 swimming further than vehicle treated animals. There was also an effect of sex (F(1,69) = 17.4, P = 0.000087) with male fish swimming further than females, similar to what we have observed previously (Rajput et al., 2022). Finally, there was a trend towards an interaction ((F(1,69) = 3.23, P = 0.077) such that males appeared to be more affected by the drug than females. FDR corrected post-hoc t-tests within sex confirmed the interaction, finding that MK-801 had no effect on distance travelled in females (P = 0.68) but did in males (P = 0.012).

**Figure 3.**
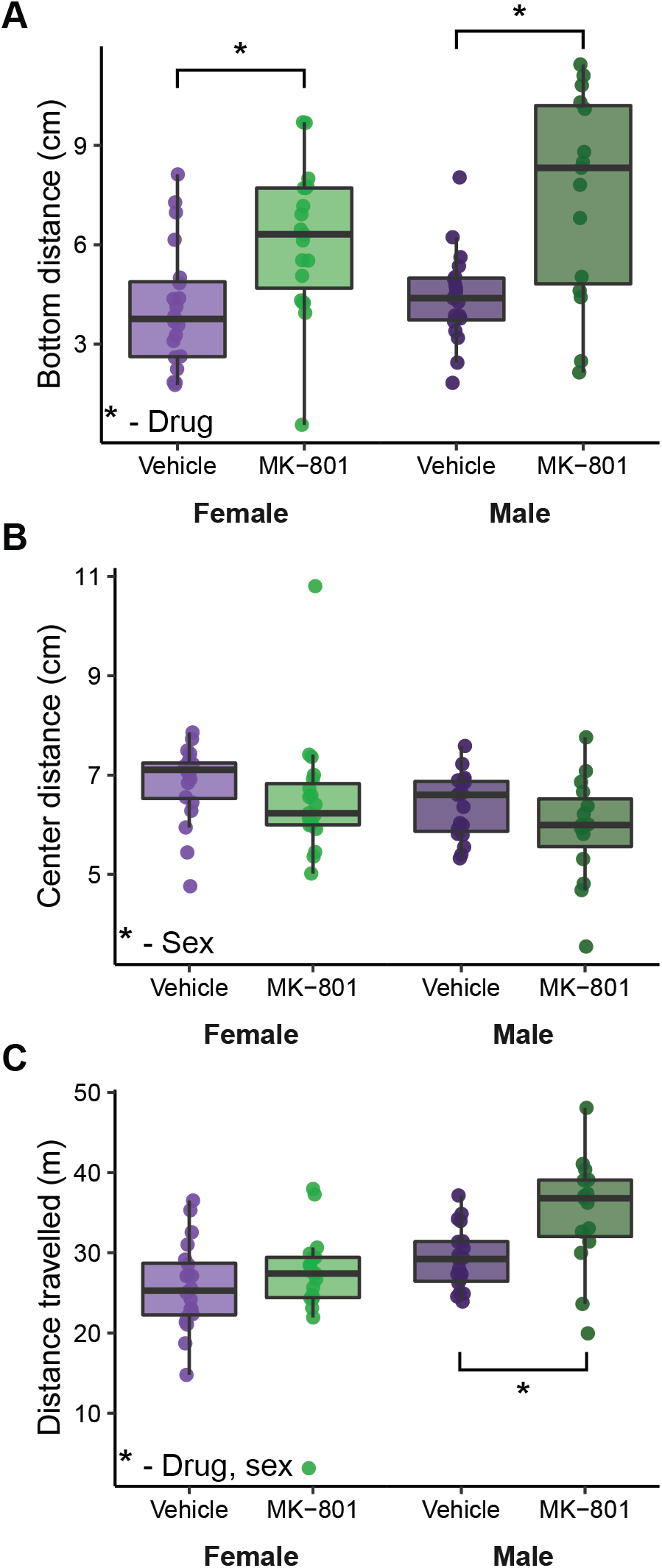
Behavioral effects of 4 mg/kg MK-801 administration 30 minutes prior to the novel tank test. We measured the effect of MK-801 on bottom distance (A), center distance (B), and distance travelled (C). Data are presented as box and whisker plots with the median (center line), interquartile range (box ends), and box ends ± 1.5 times the interquartile range (whiskers). *P < 0.05 based on FDR corrected post-hoc t-tests. Female vehicle: n=20, female MK-801: n=19, male vehicle: n = 19, male MK-801: n = 15.

## Discussion

The gelatin-based feed method we developed is a simple, precise, and non-invasive approach for drug administration to adult zebrafish. The use of gelatin, which is easily liquified, means that the feed can be made individually for each animal via pipetting, ensuring that fish are dosed according to their body weight. The addition of spirulina provides a vehicle for drug delivery that results in minimal leaching in the time frame fish eat the feed, which is typically less than 1 minute. Finally, as a proof-of-principle, we used our feed to deliver MK-801, an NMDAR antagonist, prior to the novel tank diving test. We found that MK-801 increased locomotor activity and decreased predator avoidance/anxiety-like behaviors, consistent with prior work (e.g., Menezes et al., 2015; Seibt et al., 2011; Sison and Gerlai, 2011).

Our gelatin feed provides important advantages in precision and ease of use compared to other feed-based drug delivery strategies that have been developed for adult zebrafish. For example, Sciarra and colleagues (2014) described the use of a commercial gelatin-based feed, Gelly Belly, for drug delivery. However, precise drug delivery with Gelly Belly is difficult because the food is inhomogeneous and requires cutting and weighing small amounts of solidified food for each animal. Other approaches are not easily tailored to individual fish based on weight, like a gluten-based feed described by Zang and colleagues (2011) or a gelatin/agar paste that is pressed into a 3D printed mold (Lu and Patton, 2022). These approaches administer the same amount to each fish, relying on the assumption that all animals are of approximately the same weight. However, fish vary considerably in size, so administering the same amount of feed to each animal will result in different dosing relative to body weight. For example, in the present study, the average weight of our fish was ∼250 mg with a range of 155 to 435 mg. This means a drug dose developed for the average weight would result in a 60% overdose of our smallest fish, and a 40% underdose of our largest fish.

One potential drawback to feed-based methods for drug delivery is that animals may refuse the feed if the taste of a drug is unpalatable. This can be overcome by using attractants or other additions to the feed to mask the taste. For example, additions like clam juice (Chang et al., 2019; Sciarra et al., 2014) or Power Bait, a commercial fish attractant, (Lepage et al., 2005) have been successfully used to overcome the taste of added compounds. Here, we used an extract of freeze-dried brine shrimp. Another option would be to lower the drug dose and feed fish multiple boluses to reach the appropriate dose, or increase the size of the feed (e.g., to 1.25% body weight) to reduce drug concentration.

Overall, our gelatin-based feed method provides important improvements for the administration of drugs to adult zebrafish. Most notably, our approach avoids the stress and waste associated with beaker dosing, the current most used method. Because our feed can be easily pipette into individually tailored morsels, drug delivery is based on body weight. This simple and inexpensive method brings drug administration to adult zebrafish in line with best practices for drug administration that are standard in rodent and primate research.

## Acknowledgments

This work was supported by the National Institutes of Health (R35GM142566) to JWK.

## Methods

### Subjects

Subjects were adult male and female AB and TL zebrafish 16-52 weeks of age. All fish used in experiments were bred and raised at Wayne State University. Animals were within two generations of fish originally obtained from the Zebrafish International Resource Center at the University of Oregon. Animals were kept under standard condition on high density racks (temperature 27.5 ± 0.5 °C; water conductivity 500 ± 10 μS, and a pH of 7.5 ± 0.2) with a 14:10 light/dark cycle (lights on at 8:00 AM). Fish were fed twice a day with a dry feed in the morning (Gemma 300; Skretting, Westbrook, ME, USA) and brine shrimp (*Artemia salina*; Brine Shrimp Direct, Ogden, UT, USA) in the afternoon. One week prior to behavioral testing, fish were placed as male/female pairs into 2 L tanks. Tanks were divided in half with a transparent divider with two fish in each section and a total of four fish in each tank. Body weight was recorded one day prior to experimentation by weighting fish in a beaker containing approximately 50 mL fish facility water. Fish were individually netted and gently patted twice with a dry paper towel to remove excess water prior to weighing. After experiments, animals were euthanized and sex was confirmed by the presence or absence of secondary sex characteristics (i.e., color, shape, and fin tubercles) and eggs. All procedures were approved by the Wayne State University Institutional Animal Care and Use Committee.

### Gelatin feed preparation

Our gelatin-based feed was made from a mix of gelatin, spirulina and brine shrimp extract. The brine shrimp extract was prepared by suspending 250 mg/mL of mikro fine brine shrimp (Brine Shrimp Direct) in water and stirring for one hour at room temperature. The suspension was centrifuged twice at room temperature at 12,500 g for 10 minutes, keeping the supernatant each time. Two volumes of water were then added to dilute the extract, and it was added to a tube containing spirulina (Argent Aquaculture, Redmond, WA, USA) to make a 4% w/v suspension. When drugs were added, part of the diluted extract was replaced with concentrated compound prior to mixing with spirulina to achieve the desired final concentration. The suspension was then heated at 45 °C for five minutes with periodic vortexing and added to a tube containing gelatin (Sigma-Aldrich, St. Louis, MO, USA) to make a 12% w/v mixture. We used a porcine derived gelatin with a Bloom number of ∼300 g. The mixture was then stored at − 20 °C overnight. Small morsels for feeding (at 1% body weight) were created by heating gelatin mixture to 45 °C until liquid and pipetting onto parafilm. Samples were then placed at −20 °C for at least 20 minutes to re-solidify and then kept on ice prior to feeding.

### Methylene blue leaching

We added methylene blue to determine the rate at which compound leaches from our feed. Methylene blue (Sigma-Aldrich) was added to the feed at a 2 mg/mL final concentration (equivalent to a 20 mg/kg dose). Feed samples were made at a volume of 1.75 μL as described above. Samples were placed into 1.5 mL tubes containing 50 μL water and heated to 27 °C, the same approximate temperature of our fish facility water, and left for 1, 2.5, 5, or 10 min. At each timepoint, the supernatant was removed and absorbance at 668 nm (Whang et al., 2009) was read using a NanoDrop 2000C spectrophotometer (version 1.6.198, Thermo Scientific, Waltham, MA, USA). Samples were derived from two separate preparations with 3 experimental replicates from each set. Absorbance measurements were taken in triplicate, and median values were used for analysis. Data were normalized to a sample containing the same concentration of methylene blue and brine shrimp extract used in the feed preparation for leaching, representing the maximum potential leaching.

### Gelatin-feed administration

To determine if zebrafish would eat the gelatin feed, we conducted a five-day feeding trial where our feed was given in lieu of the normal morning feed. Prior to feed administration on each day, fish were transferred from their home rack to a behavioral room and allowed to habituate for one hour. Two to five minutes prior to feeding, transparent barriers were inserted to briefly isolate fish. Feed was then given to each individual at 1% body weight. During the trial, we measured the time to eat the feed and if the feed was successfully eaten within five minutes. Barriers were removed after fish successfully ate the feed. Fish were then returned to their housing racks.

### Drug delivery and the novel tank test

As a proof of concept, we used our gelatin-based feed to deliver an NMDA receptor antagonist, (+)-MK-801 hydrogen maleate (Sigma-Aldrich), to AB fish prior to capturing their behavior in the novel tank test. For each of two days prior to drug administration, fish were fed a non-dosed gelatin feed as described above. On the day of behavioral testing, fish were transferred to the behavioral room and allowed to habituate for one hour. Feed containing MK-801 (4 mg/kg) or vehicle (water) was administered 30 min prior to behavioral testing. Animals that did not eat the feed were excluded from analysis (2 animals refused the dosed feed and 3 animals were distracted by placement of the barrier and did not eat the gelatin feed during pre-exposure days). For behavior, fish were carefully netted and placed into an open-top frosted acrylic tank (15 × 15 × 15 cm, ShopPopDisplays, Woodland Park, NJ, USA) filled with 2.5 L of fish facility water for 6 min. Water was changed between animals. The tanks were kept in a white plasticore enclosure to ensure no disturbances during video recording. Three-dimensional video recordings were captured utilizing D435 Intel Realsense™ cameras (Intel, Santa Clara, CA, USA) mounted 20 cm above the novel tanks, and fish were tracked using DeepLabCut (Mathis et al., 2018) as previously described (Rajput et al., 2022).

### Statistical analysis

Statistical analysis was done using R version 4.1.2 (R Core Team, 2016), and data were visualized using ggplot2 (Wickham, 2015). ANOVAs were performed as described. Results of behavioral experiments were followed up using false discovery rate (FDR) corrected t-tests within sex (Benjamini and Hochberg, 1995).

